# Let the prey speak: Using PNA clamps to silence predator DNA in marine faecal diet studies

**DOI:** 10.64898/2026.06.22.733645

**Authors:** Andrea M Polanowski, Leonie Suter, Bruce E Deagle, Julie C McInnes

## Abstract

DNA metabarcoding of faeces is a powerful, non-invasive method for assessing predator diets. However, when studying the diet of generalist predators, broad PCR primers are used to amplify the wide range of potential prey species and metabarcoding outputs are often dominated by sequences from the predator. While blocking primers can be used to reduce PCR amplification of predator DNA, they frequently cause partial predator suppression and unintended prey blocking. Peptide nucleic acid (PNA) clamps, offer a promising, underutilised alternative by binding strongly and selectively to predator DNA to block its PCR amplification. In this study we designed and validated a novel PNA clamp targeting the 18S rRNA gene to suppress bird and mammal predator DNA in dietary samples. We tested this clamp on tissue mixtures and faecal samples from three seabird and two seal species across temperate, subantarctic, and Antarctic regions. The PNA clamp substantially increased the proportion of prey reads recovered while maintaining consistent prey community composition across all predator species. Our results demonstrate not only the general effectiveness of PNA clamps over standard blocking primers, but also provide a powerful, broadly applicable new tool to improve the accuracy in DNA diet metabarcoding studies.

## 1 Introduction

DNA-based dietary analysis of faecal samples is a powerful, non-invasive approach for assessing diet composition, providing insights into predator diet preferences, trophic interactions and ecosystem function (Coveney et al. 2025; Ando et al. 2020). This approach has been widely applied across diverse taxa, from apex predators to invertebrates and herbivores (see Table S1 in Convey et al. 2025; Ando et al. 2020). When using DNA metabarcoding to study generalist predator’s diet, universal eukaryotic primers are ideal because they allow simultaneous detection of a taxonomically broad range of potential food. An issue when using such highly conserved primers is that predator DNA is co-amplified and dominates sequencing outputs (Cuff et al. 2023). For example, in a diet study of green sea turtles up to 99.8% of recovered sequences came from the turtle (Díaz-Abad et al. 2022). The food DNA component can be increased by optimising collection protocols, such as selecting fresh samples, or avoiding fasting individuals, but predator DNA consistently remains dominant in recovered sequences (McInnes et al. 2017a).

The most straightforward approach to overcome the low amount of food DNA is by increasing sequencing depth. However, this approach is inefficient and costly; in a copepod diet study prey sequences accounted for only 0.4% to 1.2% of the data, requiring extremely high sequencing depth to characterise diet (Flo et al. 2023). This approach has several other limitations: required sequencing depth is difficult to predict without pilot studies, prey detection varies among samples, and bioinformatic processing becomes more complicated. An alternative approach is the use of group-specific primers, which exclude predator DNA but reduce taxonomic breadth (Cuff et al. 2023).

Designing such primers can be challenging in the absence of comprehensive reference databases, and may result in unintended biases, including failure to detect poorly characterised or unexpected prey taxa (Coveney et al. 2025; Cuff et al. 2023). For generalist predators, recovering the full dietary breadth requires the use of multiple different marker sets, and linking the resulting datasets to obtain a picture of overall diet is problematic (Deagle et al. 2019).

Blocking primers are a widely used method to supress predator DNA. These modified oligonucleotides bind preferentially to predator sequences but are chemically modified to prevent amplification (Vestheim & Jarman, 2008). Blocking primers have been widely applied in marine predator diet studies, including penguins (McInnes et al. 2016; Jarman et al. 2013), seals (Jimbo et al. 2021; Thomas et al. 2022; Deagle et al. 2009) and whales (Carroll et al. 2019; De Vos et al. 2018). However, blocking primers are variable in their effectiveness to block predator DNA amplification (Homma et al. 2022), and can inadvertently inhibit amplification of closely related prey (Zabala et al. 2025; Piñol et al. 2015; Jarman et al. 2013), as they are usually most effective when they overlap PCR primer binding sites which tend to have low variability between species for universal primers (Vestheim & Jarman, 2008).

PNA clamps offer a promising alternative. These synthetic DNA analogues enable stronger binding than regular DNA and can target taxon-specific variable regions between PCR primers, allowing them to physically block PCR amplification of target taxa (https://pnabio.com/pcr-blockers). This method routinely reduces contamination in plant microbiome studies (Hussain et al. 2025; Mertin et al. 2025; Flörl & Bokulich, 2025) and detects rare cancer mutations in medical diagnostics (Chen et al. 2023; Fouz & Appella, 2020), yet remains underutilised in dietary metabarcoding. They have successfully improved prey detection in aquatic taxa including lobster larvae (Ascher et al. 2025), krill (Cleary et al. 2012) and several fish (Homma et al. 2022; Sakaguchi et al. 2017; Terahara et al. 2011), with Homma et al. (2022) reporting 99% host suppression compared to just 33% using traditional blocking primers. This underutilization is highlighted in a review by Coveney et al. (2025) of 85 marine vertebrate metabarcoding dietary studies (2005 to 2023), which found that while 30 studies used blocking primers, only three utilized PNA clamps. Their application to seabirds and marine mammals is limited, with only a single study in Bryde’s whales (Carroll et al. 2019).

In this study, we design a PNA clamp for a widely adopted universal 18S primer set to suppress predator DNA amplification in seabird and marine mammal dietary studies. This universal marker targets a broad range of diet taxa, but sequences recovered are often dominated by predator DNA (Satgé et al. 2024; Kennerley et al. 2024; Fayet et al. 2021; Cavallo et al. 2020; Ratcliffe et al. 2021; McInnes et al. 2017a, 2017b). This study aims to 1) evaluate the PNA clamp using tissue mixtures and faecal samples collected across temperate, subantarctic, and Antarctic regions, 2) assess its ability to reduce predator read proportions and increase prey detection and 3) evaluate whether the clamp alters the relative representation of prey taxa. The results show the PNA clamp will be widely useful to improving dietary resolution for generalist marine predators.

## 2 Methods

### 2.1 PNA clamp and primers

A PNA clamp was designed to block amplification of bird and mammal DNA for the V7 region of the 18S rRNA gene. Predator reference sequences were aligned with potential prey sequences and submitted to PNA Bio (www.pnabio.com) for clamp design. Initial PCR optimisation indicated optimal clamp annealing temperature of 75°C and PNA clamp concentrations ranging from 2.5 µM to 5 µM (Supplementary Methods S1).

For PNA concentration optimisation (see section 2.3) we used primers 18S_SSU_F _a and 18S_SSU_R (Table 1) which were originally designed for use with a blocking primer. For the PNA testing on faeces and tissue (see section 2.4) we utilised an established 18s primer set, 18S_SSU_F primer paired with the same reverse primer as above, which has previously been validated in faecal diet studies (Cavallo et al. 2020; Ratcliffe et al. 2021; McInnes et al. 2017a). Both primer pair combinations target an overlapping 18S region and work with the same PNA clamp (supplementary Figure S1).

**Table 1.**
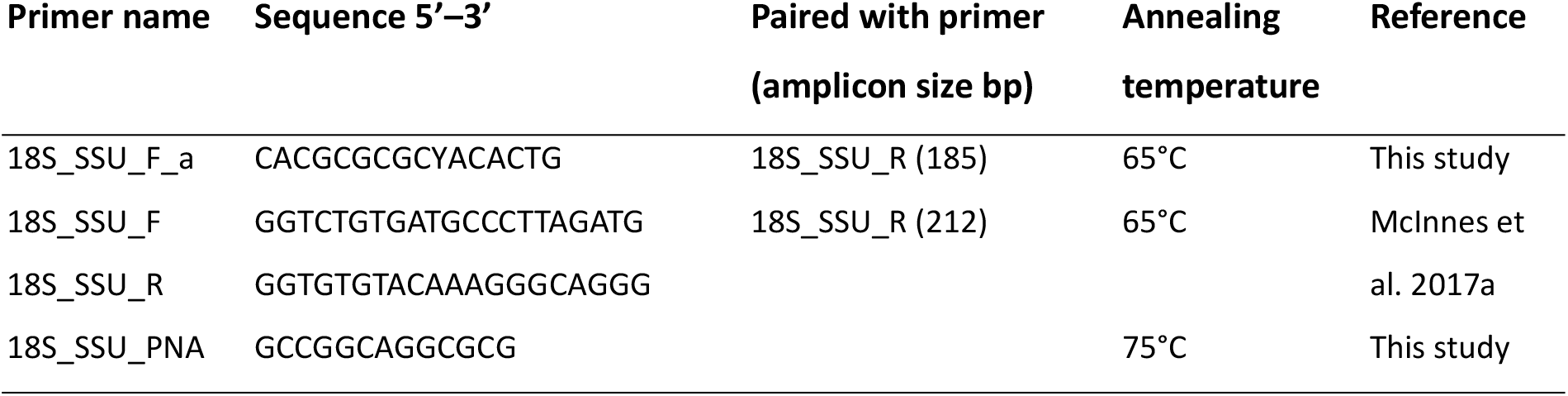
Oligonucleotides and PNA clamp used in this study.

### 2.2 Samples

For the tissue comparison, DNA was extracted from two predators - bird (shy albatross, *Thalassarche cauta*) and mammal (mouse, *Mus musculus*) – and three prey types: fish (Antarctic lanternfish, *Electrona antarctica*), krill (Antarctic krill, *Euphausia superba*) and squid (arrow squid, *Nototodarus gouldi*). For the diet study, faecal samples were collected from three seabird species (gentoo penguin, *Pygoscelis papua*; shy albatross, *T. cauta*; and black-browed albatross, *T. melanophris*) and two seal species (Australian fur seal, *Arctocephalus pusillus doriferus* and Antarctic fur seal, *A*. gazella; see supplementary Table S1 for sample information).

### 2.3 PNA concentration optimisation

To determine the lowest effective PNA clamp concentration (2.5 µM or 5 µM) we conducted trials using both tissue mixes (n=4; see supplementary Figure S2) and faecal samples including gentoo penguins (n=14), shy and black-browed albatrosses (n=4 of each) and Australian fur seals (n=2).

PCRs were performed in two rounds (Suter et al. 2021) using three PNA concentrations (no clamp, 2.5 µM, and 5 µM) with a positive control (squid; *Psychroteuthis sp*.) and a No Template Control (NTC). Round one amplified the target gene using 0.2 µM of primers (18S_SSU_F_a and 18S_SSU_R; Table 1), 8 bp MIDs and Illumina sequencing primers. The 10 μL mix contained 5 μL AmpliTaq Gold™ 360 (Life Technologies), 1 x EvaGreen (Biotium), 0.2 μL of bovine serum albumin and 1 μL DNA template. Cycling conditions were 95°C (10 min); 37 cycles of 95°C (30 sec), 75°C (10 sec, PNA clamping), 65°C (30 sec), 72°C (30 sec), and 72°C (7 min) on a LightCycler 480 (Roche Diagnostics). Round one PCR products were diluted 1:10. Round two PCRs (10 µL) included 5 µL AmpliTaq Gold™ 360 , 0.1 µM each of Illumina adapters with 8bp MIDs, and 2 µL of diluted template for 10 cycles: 95°C (30 sec), 55°C (20 sec), 72°C (45 sec); and 72°C (5 min).

PCR products were pooled and purified (0.8x) with Agencourt AMPure XP beads (Beckman Coulter, Brea, CA, USA). The fragment size of the library was checked on a 2100 DNA bioanalyzer (Agilent Technologies, Santa Clara, CA, USA), and DNA was quantified on a QUBIT 2.0 fluorometer (Life Technologies, Carlsbad, CA, USA) and diluted to 4 nM. Sequencing was performed at the Ramaciotti Centre for Genomics (UNSW, Sydney, Australia) on an Illumina MiSeq platform (150 bp paired-end).

### 2.4 PNA testing on tissue and faeces

To evaluate the effect of the PNA clamp on dietary read proportions, we compared DNA tissue mixtures and faecal samples with either no clamp or a 2.5 µM concentration. Tissue mixes of equal proportion were tested in triplicate across four groups: 1) individual species, 2) single predator with single prey, 3) combined predators, and 4) all five species combined (see supplementary Table S2 for tissue mixtures). Additionally, we tested the clamp on 24 faecal samples each from five predator species: gentoo penguins, shy and black-browed albatrosses, Australian and Antarctic fur seals. Round one PCRs matched the optimisation protocol (section 2.3), except with primers (18S_SSU_F &_R) with either no clamp or a 2.5 µM concentration. Controls, cycling, round-two PCRs, and sequencing followed the same workflow.

Bioinformatics processing and analysis followed the methods of Suter et al. (2021) and a brief description can be found in Supplementary Methods S2. Taxonomic assignments were grouped into four categories: predator (bird or mamma), prey (fish, krill or squid), parasite (Conoidasida or Cestoda) or “other – non diet”. Reads were summarised using plyr (Wickham, 2011) and visualised with ggplot2 (Wickham, 2016). Paired Wilcoxon signed-rank tests, with Holm-adjusted p-values, evaluated how the PNA clamp affected diet read proportions (total diet reads / total reads per sample) for each predator. A Pearson correlation was calculated to assess the effect on diet read proportion (proportion of diet item reads relative to total diet reads) within each sample with and without using of the PNA clamp.

## 3 Results

### 3.1 PNA Concentration

A clamp concentration of 2.5 µM inhibited predator DNA amplification as effectively as 5 µM in gentoo penguin scats (Figure 1A), and across three additional predator species and tissue DNA (see supplementary Figure S2); supporting the choice of 2.5 µM as the optimal concentration for reagent efficiency.

**Figure 1.**
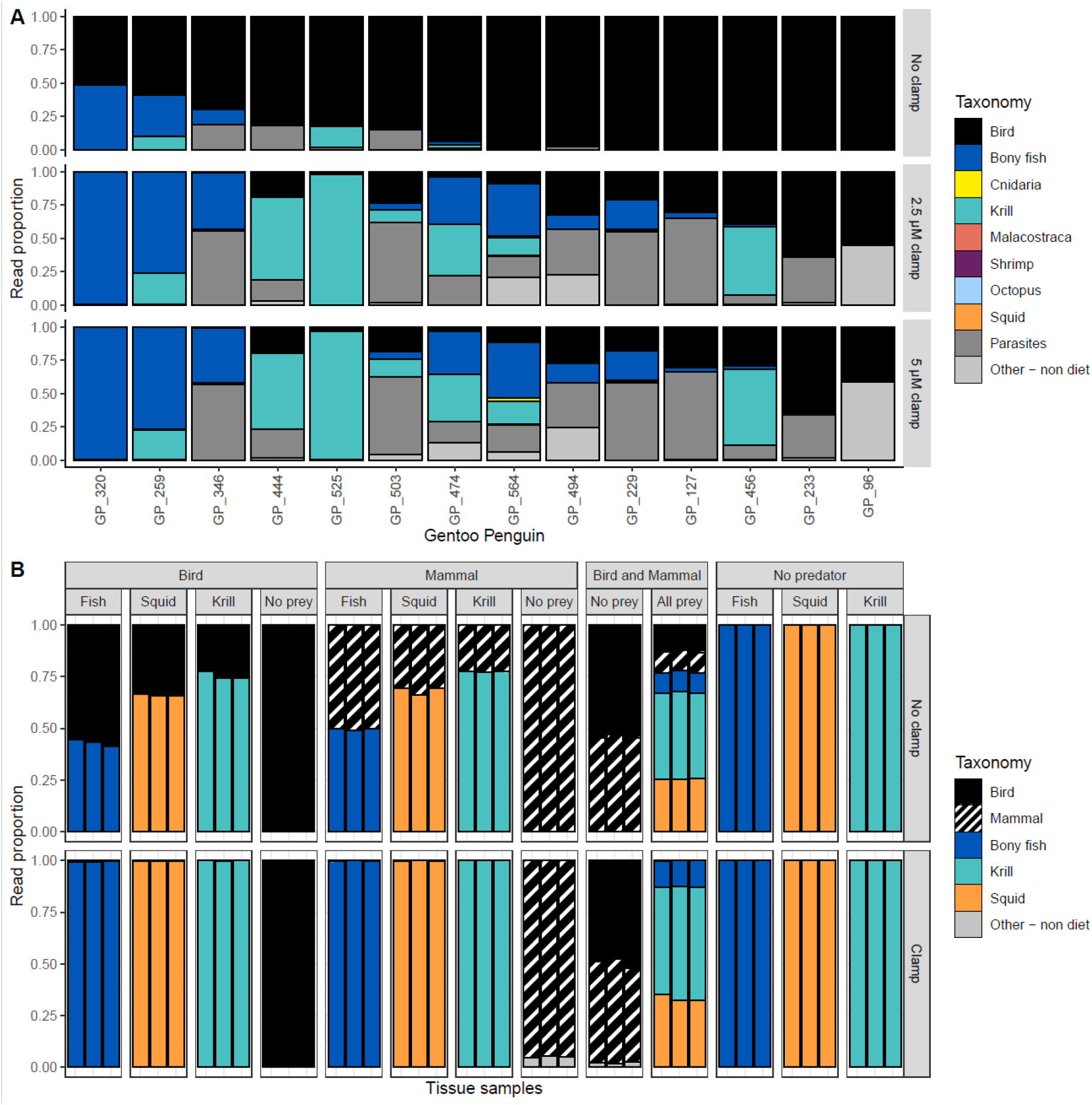
(A) Taxonomic read proportions for Gentoo penguin faecal samples (n=14) across three PNA clamp concentrations (no clamp, 2.5 µM, and 5 µM). (B) Taxonomic read proportions for tissue-based optimization of the PNA clamp with four experimental treatments; 1) individual species, 2) single predator with single prey, 3) both predators combined, and 4) all five species combined with either no clamp or a 2.5 µM concentration.

### 3.2 PNA testing on tissue

Using tissue mixes, the PNA clamp demonstrated high specificity, effectively suppressing predator DNA across both bird and mammal samples where prey DNA was present, without inhibiting the amplification of diverse prey groups (fish, squid, and krill). Where no prey was present, predator DNA was amplified with and without PNA clamp (Figure 1B).

### 3.3 PNA testing on faeces

Across all predator species, the PNA clamp improved prey detection in faecal samples compared to no clamp (see Figure 2 for gentoo penguin and Australian fur seals, and Figure S3 for remaining three predators). This included samples with low or previously undetected prey concentrations (e.g., GP_525, GP_108, AuFS_025, AuFS_340). Across all samples, the use of the PNA clamp reduced predator DNA read proportions from 60% without PNA clamps to 7% with PNA clamps, while prey DNA read proportions increased from 38% without PNA clamp to 69% with PNA clamp. Prey DNA increased proportionally to the other DNA sources, with increases in non-prey taxa such as parasites. This was particularly evident in gentoo and albatross samples that contained a lot of parasite DNA (Figure 2A, Figure S3).

**Figure 2.**
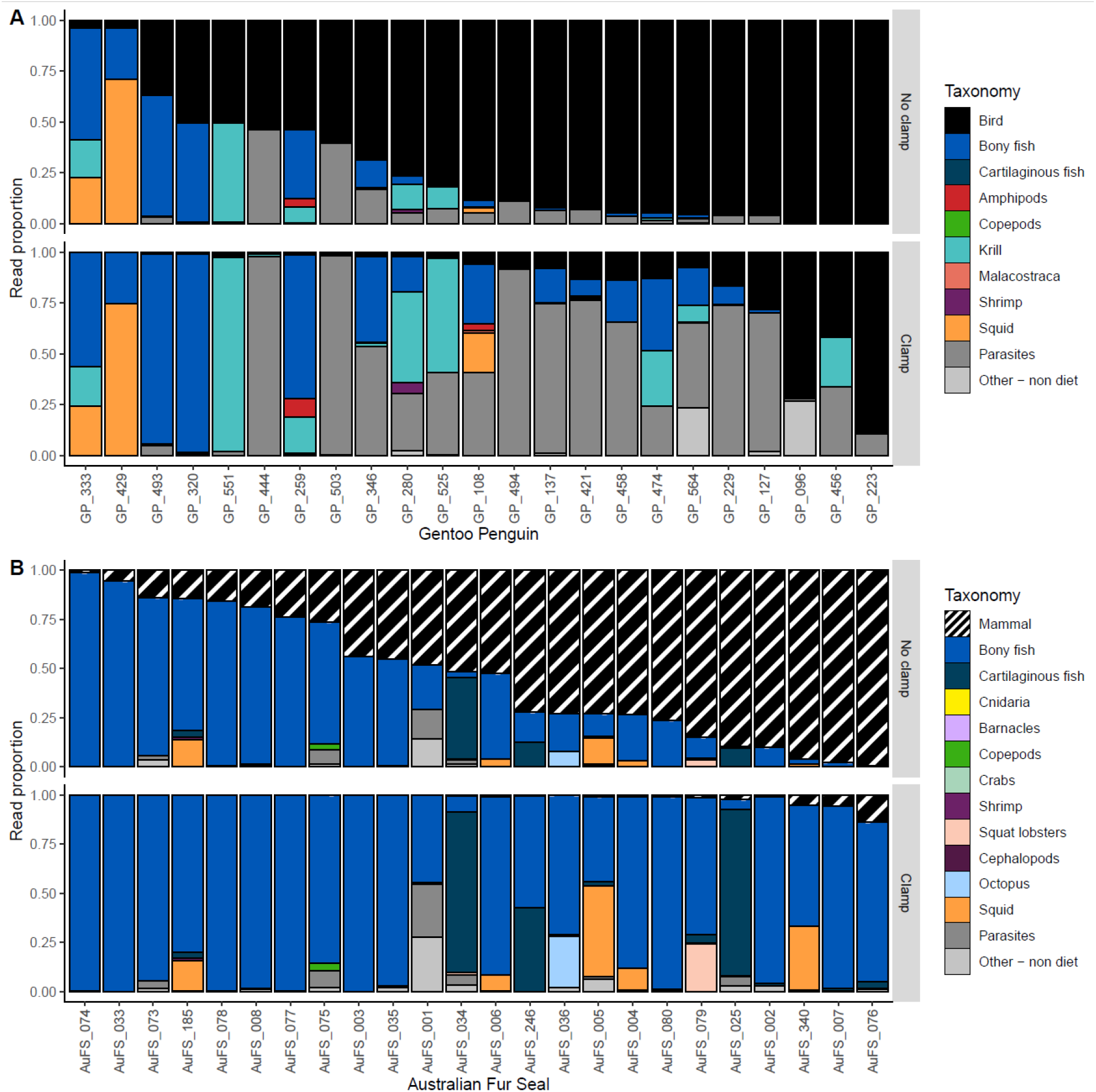
Taxonomic read proportions for (A) gentoo penguin and (B) Australian fur seal faecal samples across two PNA clamp concentrations (no clamp and 2.5 µM).

The PNA clamp significantly increased the detected prey proportion across all five predator species compared to the no-clamp control (p < 0.001 for all species, Figure 3; supplementary Table S3). Diet reads increased between 21% (black-browed albatross, lowest increase) to 50% (Australian fur seal, highest increase).

**Figure 3.**
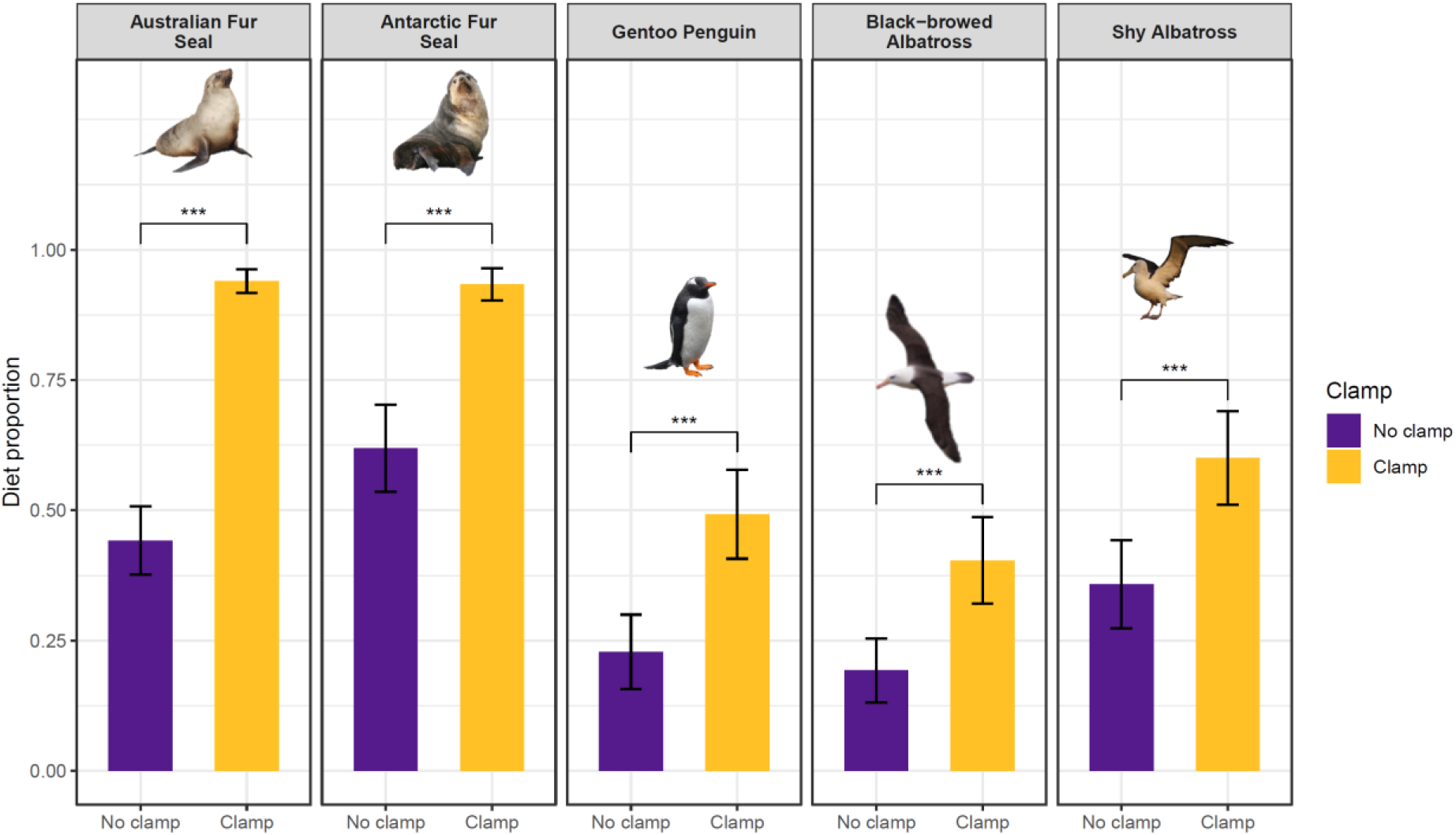
Comparison of prey DNA proportions detected with and without a PNA clamp across five marine predator species indicate that the use of PNA clamp significantly increased the proportion of diet reads across all five predators. Asterisks indicate adjusted p-values of paired t-tests (***: p-adjust < 0.001).

The PNA clamp did not affect the proportion of reads assigned to each detected diet item: when plotting diet proportion (reads of one diet item/total diet reads per sample) detected without clamp against diet proportion detected with clamp, a nearly perfect correlation (*R* = 0.96, *df* = 196, *p* <0.001) was found across all five predators (Figure 4). The few samples that deviated from this relationship had very few diet reads recovered, indicating that low target DNA in a sample may introduce PCR stochasticity and add noise to the data.

**Figure 4.**
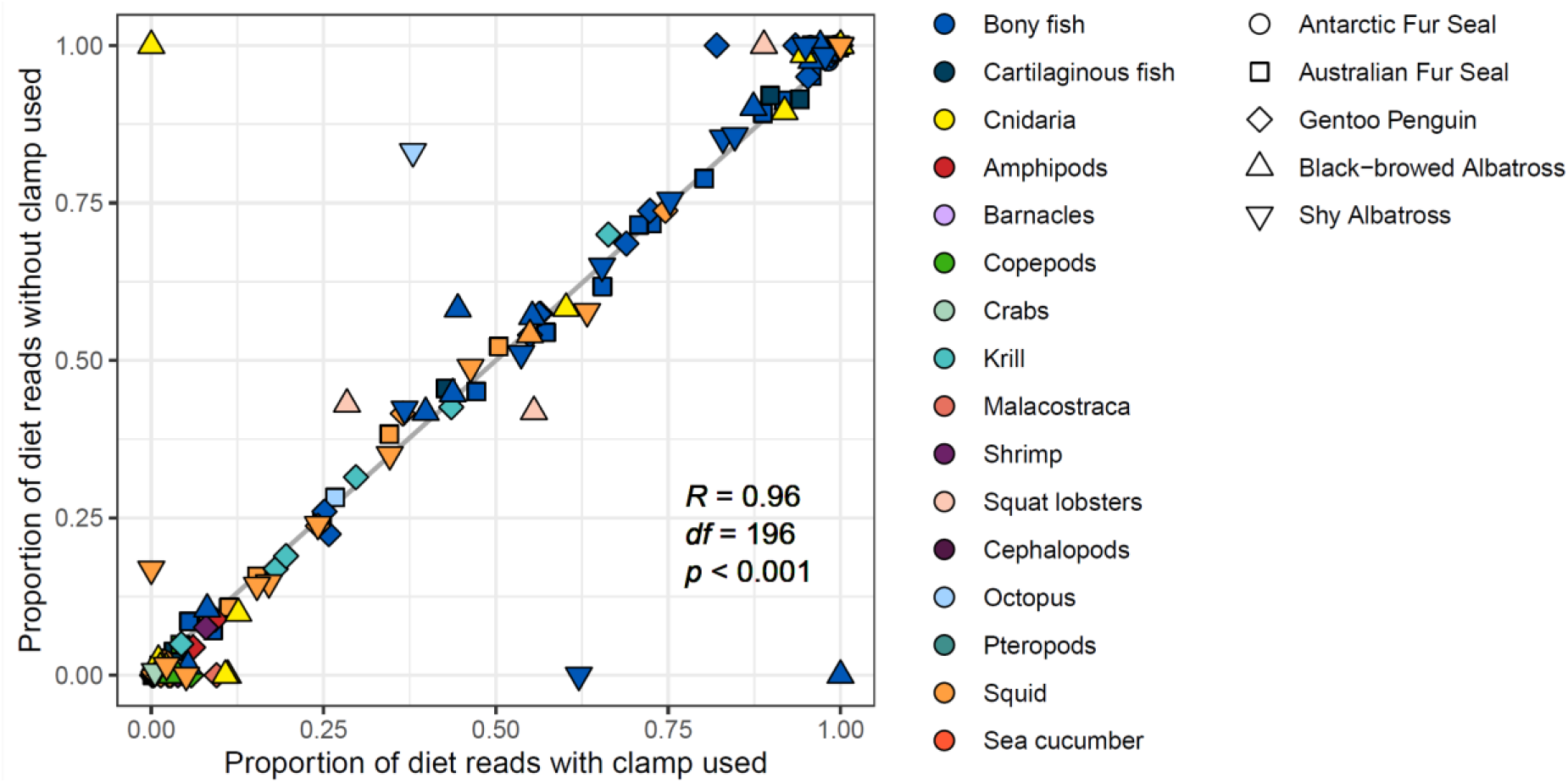
Proportion of diet reads (reads of one diet item / total diet reads) detected from faecal samples of five predators, with the use of PNA clamp (x axis) and without the use of PNA clamp (y axis).

## 4 Discussion

Our results demonstrate that our PNA clamps effectively reduce predator DNA amplification and substantially increase prey read proportions in metabarcoding studies of seabird and marine mammal diets. Compared with blocking primers, which may partially block prey or fail to completely exclude predator DNA, this PNA clamps provide a highly specific and effective alternative. These findings align with previous research (Ascher et al. 2025, Homma et al. 2022; Terahara et al. 2011) showing near-complete suppression of predator DNA and demonstrate that, when optimised, PNA clamps are a robust tool for broadening taxonomic resolution in marine predator diet studies.

Importantly, the PNA clamp significantly reduced the proportion of predator DNA and increased prey detection without biasing prey composition. This is a marked improvement on traditional blocking primers. The only instances where the prey proportions showed a considerable difference between samples amplified with and without a clamp was when the initial prey proportion was very low. This was likely a result of PCR stochasticity adding noise to the data. The inclusion of samples with very low diet reads should be considered with caution as the results may not be representative or meaningful.

The use of PNA clamps also represents a cost-effective alternative to increasing sequencing depth. Without a clamp, predator DNA can exceed 95% of reads, requiring a two to 10-fold increase in sequencing depth to recover sufficient prey data. Although sequencing costs have decreased, this still represents a significant expense, and larger data volume can be difficult to process bioinformatically. In contrast, PNA clamps can substantially improve read allocation at minimal cost (∼AUD $0.64 per sample). The 18S clamp used in this study reduces predator DNA by an average of 53%, requiring lower sequencing depth per sample and therefore allowing more samples in a single MiSeq run.

As with all universal makers, there is still the need to have some a priori understanding of the animal diet prior to assessing the value of the bird and mammal clamp. The clamp will block prey DNA if the predator consumes bird or mammal in their diet, e.g. skuas, giant petrels and leopard seals (to name a few). These species are problematic for dietary studies using universal 18S markers as there is no way to differentiate whether recovered bird or mammal DNA originated from the predator or prey. In such cases group specific markers would be needed with either considerable sequencing depth, or predator-specific PNA clamps, e.g. a skua specific clamp used with a marker where the amplicon varies between bird taxa.

The increase in prey DNA is proportional to other non-prey DNA in the samples, irrespective of the reduction in predator DNA. This means that if a sample was dominated by parasites or environmental contamination (e.g. sampling substrate), then the prey sequence reads will still remain relatively low even with the use of the clamp. For example, some of the albatross samples in our study were collected from birds that were fasting during incubation or collected off a dirt substrate (increasing the detection of vegetation or fungi DNA). In these samples, the PNA clamp removed most of the predator DNA, yet sequences were dominated by non-prey sources (e.g. parasites or fungi; Figure S3), as simply not enough prey DNA was present in these samples. This shows the importance of good collection protocols and understanding the biology of the predator (McInnes et al. 2017a; Oehm et al. 2011). However, these data highlight the value of the clamp not only for determining diet, but also for assessing parasite loads. The combination of the PNA clamp used to reduce predator DNA as seen in this study, combined with sample collections during fasting which reduces prey DNA (McInnes et al. 2017a), is likely to be a winning solution for parasitologists.

Overall, this PNA clamp offers an efficient, robust and cost-effective method to suppress predator DNA and enhance prey detection in DNA metabarcoding studies. Its high specificity and efficiency make it a valuable tool for advancing ecological diet analysis in seabird and marine mammals.

## Author contribution

Bruce Deagle, Andrea Polanowski and Julie McInnes conceived the ideas and designed methodology; Andrea Polanowski and Julie McInnes collected the data; Leonie Suter and Julie McInnes analysed the data; Andrea Polanowski and Julie McInnes led the writing of the manuscript. All authors contributed to the drafts and gave final approval for publication.

## Acknowledgments

We thank Norm Ratcliffe and Kieran Love (British Antarctic Survey) for providing the gentoo scat samples. This research was funded by the Australian Antarctic Science Program: AAS 4556 and 4538.

## Data availability

Data generated during this study are available from the Australian Antarctic Division Data Centre (https://data.aad.gov.au/) with a DOI provided on acceptance.

