## Supplementary Information for "Let the prey speak: Using PNA clamps to silence predator DNA in marine faecal diet studies"

**Supplementary Information Contents**

**Supplementary Methods S1**. Initial PNA clamp optimisation.

**Supplementary Methods S2**. Bioinformatics and Analysis.

**Table S1**. Faecal samples used in this study.

**Table S2**. Composition of tissue mixtures with known predator-to-prey DNA ratios used for PNA testing. Shaded cells indicate the species included in each blend as individual samples or equal-part mixtures. All mixtures were analysed in triplicate.

**Table S3**. Effect of using the PNA clamp on diet proportion of five predator species, calculated with a paired Wilcoxon signed-rank test.

**Figure S1**. A 212 bp region of king penguin (*Aptenodytes patagonicus*, Accession Number: XR_012997719.1) 18S ribosomal RNA sequence showing the primer pairs and PNA clamp used in this study (°C is annealing temperature).

**Figure S2**. Proportion of taxonomic reads across three PNA clamp concentrations (no clamp, 2.5 µM, and 5 µM) for faecal samples of: (A) shy albatross (*Thalassarche cauta*), n = 4; (B) Australian fur seal (*Arctocephalus pusillus doriferus*), n = 2; (C) black-browed albatross (*Thalassarche melanophris*), n = 4; and (D) four experimental tissue mixes, which included equal amounts of the following DNA combinations: bird-fish, bird-krill, bird-fish-krill-mammal and bird-mammal.

**Figure S3**. Taxonomic read proportions for 24 faecal samples across two PNA clamp concentrations (no clamp and 2.5 µM). Results are shown for: (A) shy albatross (*Thalassarche cauta*), (B) black-browed albatross (*Thalassarche melanophris*), and (C) Antarctic fur seal (*Arctocephalus gazella*).

Supplementary Methods S1. Initial PNA clamp optimisation.

To optimise the PNA clamp annealing temperature, PCRs were performed using shy albatross (*Thalassarche cauta*) tissue, a gentoo penguin (*Pygoscelis papua*) faecal sample and a No Template Control (NTC). Trials were conducted across a 68–78°C temperature gradient using four PNA concentrations: no clamp, 1.0 µM, 2.5 µM and 5.0 µM. The PCR mix (10 µL) contained 5 µL AmpliTaq Gold™ 360 Master Mix (Life Technologies), 0.2 µM for each forward (18S_SSU_F_a; CACGCGCGCYACACTG) and reverse (18S_SSU_R; GGTGTGTACAAAGGGCAGGG) primer, 1 x EvaGreen (Biotium), 0.2 µL of bovine serum albumin and 1 µL DNA template. Thermal cycling conditions were 95°C for 10 min, followed by 37 cycles of 95°C for 30 sec, 68-78°C for 10 sec (PNA clamping), 65°C for 30 sec, and 72°C for 30 sec, with a final extension of 72°C for 7 min on a LightCycler 480 (Roche Diagnostics). The samples were loaded on the Agilent DNA bioanalyzer (Agilent Technologies, Santa Clara, CA, USA) and the concentration of the desired band was compared for inhibitory effects. The optimal annealing temperature was 75°C and a PNA clamp concentration between 2.5 µM and 5 µM.

Supplementary Methods S2. Bioinformatics and Analysis.

Sequencing reads were processed using a USEARCH pipeline (v11.0.667; Edgar, 2010; Suter et al. 2021). In brief, paired reads were merged (fastq_mergepairs), and first round MID sequence pairs were filtered and trimmed using R package ShortRead (Morgan et al. 2009). Sequences were dereplicated (fastx_uniqes), zOTUs identified (unoise3), and a zOTU table was calculated (otutab). Additional chimaeras were removed using dada2 (Callahan et al. 2016). The resulting zOTUs were compared to the NCBI nucleotide database, excluding environmental sequences. Taxonomy was assigned using MEGAN (Huson et al. 2016). Read values smaller than 15 were set to zero to remove noise. No contamination of target taxa was detected in controls.

Table S1. Faecal samples used in this study.

| Species | Number of samples included | | Region | Location | Latitude/Longitude |
|---|---|---|---|---|---|
| | PNA concentration experiment | PNA testing on tissue/ faeces experiment | | | |
| Shy albatross | 4 | 24 | Temperate | Albatross Island, Tasmania, Australia | -40.377, 144.655 |
| Black-browed albatross | 4 | 24 | Subantarctic | Macquarie Island, Tasmania, Australia | -54.229, 158.892 |
| Gentoo penguin | 14 | 24 | Antarctic | South Georgia, Southwest Atlantic Ocean | -54.251, -36.498 |
| Australian fur seal | 2 | 24 | Temperate | Tasmania, Australia | -43.383, 146.993 |
| Antarctic fur seal | 0 | 24 | Subantarctic | Macquarie Island, Tasmania, Australia | -54.229, 158.892 |

Table S2**.** Composition of tissue mixtures with known predator-to-prey DNA ratios used for PNA testing. Shaded cells indicate the species included in each blend as individual samples or equal-part mixtures. All mixtures were analysed in triplicate.

| | | Species | Tissue mixes | | | | | | | | | | | | |
|---|---|---|---|---|---|---|---|---|---|---|---|---|---|---|---|
| Predator | Bird | *Thalassarche cauta* | ■ | | | | | ■ | ■ | ■ | ■ | | | | ■ |
| | Mammal | *Mus musculus* | | ■ | | | | | | | ■ | ■ | ■ | ■ | ■ |
| Prey | Fish | *Electrona antarctica* | | | ■ | | | ■ | | | | ■ | | | ■ |
| | Krill | *Euphausia superba* | | | | ■ | | | ■ | | | | ■ | | ■ |
| | Squid | *Nototodarus gouldi* | | | | | ■ | | | ■ | | | | ■ | ■ |

Table S3: Effect of using the PNA clamp on diet proportion of five predator species, calculated with a paired Wilcoxon signed-rank test.

| **Predator Species** | **N** | **Median (No Clamp)** | **Median (Clamp)** | **V** | **p-value** | **Adj. p-value (Holm)** | **Significance** |
|---|---|---|---|---|---|---|---|
| Antarctic Fur Seal | 22 | 0.871 | 0.989 | 0 | <0.001 | <0.001 | *** |
| Australian Fur Seal | 24 | 0.363 | 0.981 | 0 | <0.001 | <0.001 | *** |
| Gentoo Penguin | 20 | 0.051 | 0.489 | 0 | <0.001 | <0.001 | *** |
| Black-browed Albatross | 20 | 0.071 | 0.299 | 1 | <0.001 | <0.001 | *** |
| Shy Albatross | 22 | 0.126 | 0.826 | 0 | <0.001 | <0.001 | *** |

Figure S1. A 212 bp region of king penguin (*Aptenodytes patagonicus*, Acc. No. XR_012997719.1) 18S ribosomal RNA sequence showing the primer pairs and PNA clamp used in this study (°C is annealing temperature).

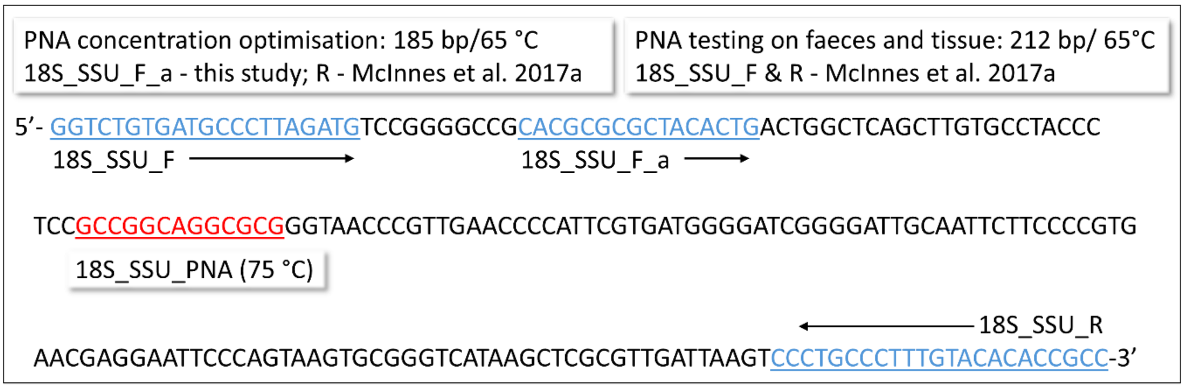


Figure S2. Proportion of taxonomic reads across three PNA clamp concentrations (no clamp, 2.5 µM, and 5 µM) for faecal samples of: (A) shy albatross (*Thalassarche cauta*), n = 4; (B) Australian fur seal (*Arctocephalus pusillus doriferus*), n = 2; (C) black-browed albatross (*Thalassarche melanophris*), n = 4; and (D) four experimental tissue mixes, which included equal amounts of the following DNA combinations: bird-fish, bird-krill, bird-fish-krill-mammal and bird-mammal.

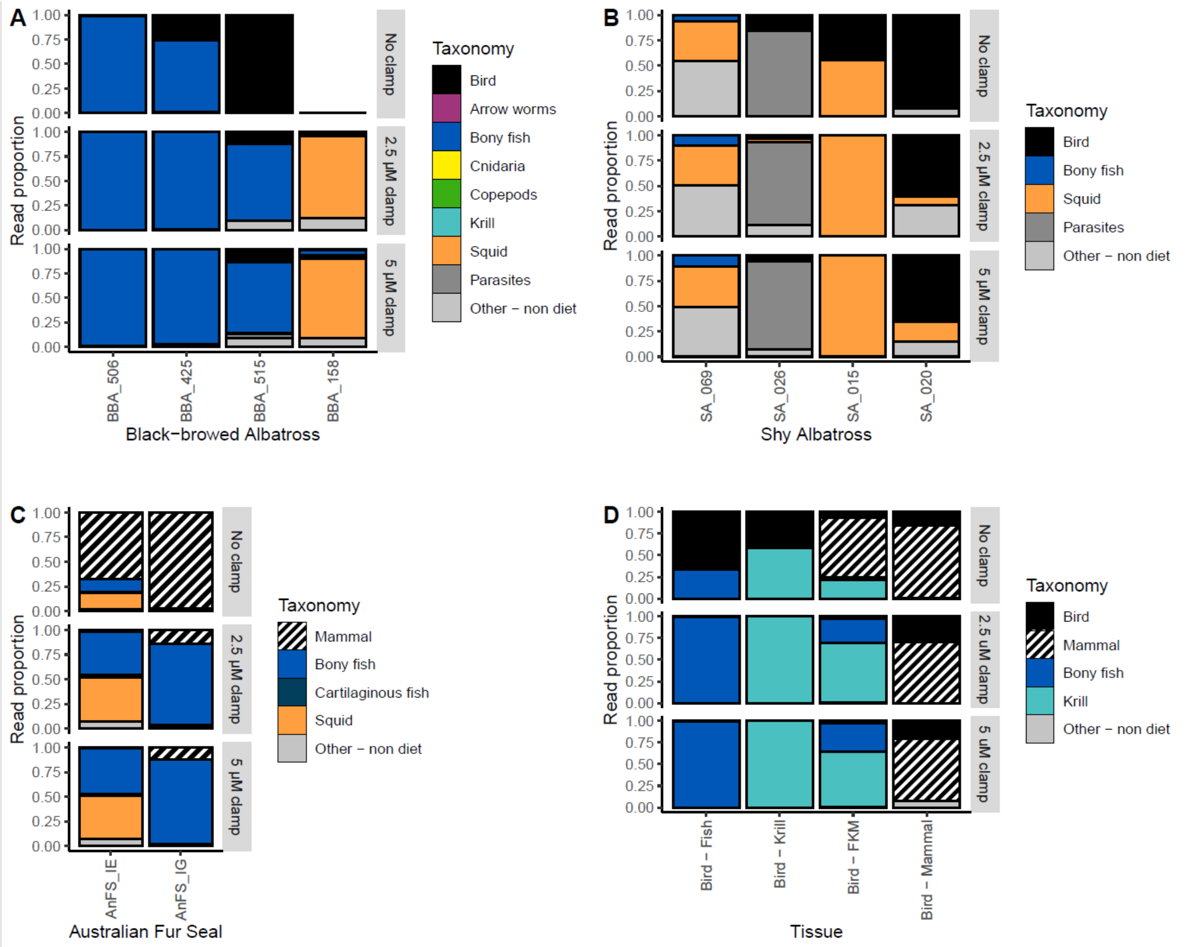

Figure S3. Taxonomic read proportions for 24 faecal samples across two PNA clamp concentrations (no clamp and 2.5 µM). Results are shown for: (A) shy albatross (*Thalassarche cauta*), (B) black-browed albatross (*Thalassarche melanophris*), and (C) Antarctic fur seal (*Arctocephalus gazella*).

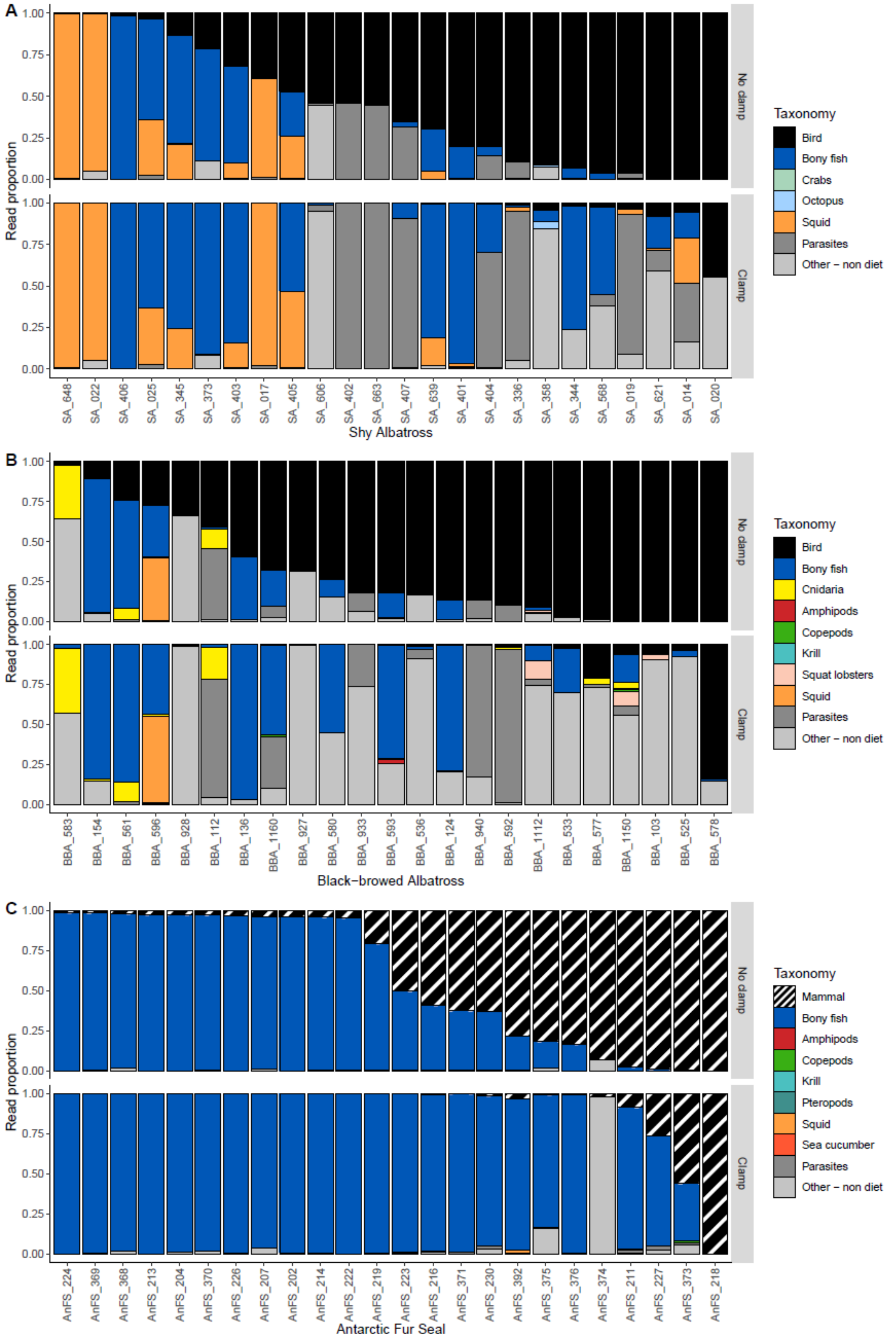
